# An electronically steerable epidural ultrasound interface for deep brain neuromodulation in freely moving rats

**DOI:** 10.64898/2026.09.10.750612

**Authors:** L Ratz, H Rivandi, GK Wardhana, E Sarkar, M Aqamolaei, Y Tong, S Desmarais, L Sohail, PD Donaldson, V Aloia, M Arlotti, VA Coenen, DG Muratore, GD Spyropoulos, TLM da Costa, MD Döbrössy

## Abstract

Low-intensity focused ultrasound (LI-FUS) clinical trials exploit either the neural activity modulating, or the blood-brain barrier opening, capacity of this stimulation modality. However, LI-FUS currently is applied only transcranially which means that it is conducive only for episodic and intermittent stimulation, although, clinical data shows that in numerous neurological and psychiatric applications, chronic and continuous stimulation is required for long-term, stable therapeutic effect.

The paper presents experimental data on a novel and innovative approach describing an implantable, epidural focus ultrasound (eFUS) device, designed for continuous, chronic and multi-site steerable neuromodulation. The miniaturized eFUS device consists of a two-dimensional piezoelectric transducer array directly integrated onto a custom ASIC, specifically engineered for proof-of-principle neuromodulation studies in the rat brain. The system generates electronically steerable focused ultrasound with software-defined focal coordinates, sufficient to stimulate neuronal activity in deep brain structures. *In vitro* acoustic characterization confirmed accurate beam steering and focusing, while *in vivo* validation demonstrated reliable stimulation of a deep subcortical target with measurable physiological effects. eFUS-mediated targeting of the ventral tegmental area in awake and freely moving rats produced increase in dopamine release in the nucleus accumbens as confirmed using fiber photometry recordings. Post-mortem histological analysis of the target regions showed the absence of inflammatory markers, although the epidural placement of the eFUS device was associated with mild tissue damage. Overall, the study provides in vitro data demonstrating the energy efficiency, and steerability of the technology, and in vivo physiological evidence of neuromodulatory ability of a deep, subcortical brain structure.

## Introduction

Neuromodulation refers to chemical, and more commonly technology-mediated, interaction with the peripheral or central nervous system to achieve therapeutic effects. It is a powerful and fast-growing treatment strategy used with patients with brain disorders – both neurological or psychiatric – which can bring long-term relief via both identified and yet unknown mechanisms modulating inhibitory and excitatory pathways, and ultimately, regional network activities ^1,2^. Over the past three decades, cross disciplinary teams including clinicians, pre-clinical researchers, biomedical, material, and computer engineers have developed and improved devices that have either become routine and FDA-approved or are designated as experimental treatments in clinical trials, testing their safety and functional efficacy as symptomatic management of brain disorders^3^.

Amongst the most frequently used neuromodulatory approaches today are Transcranial Magnetic Stimulation (TMS) and Deep Brain Stimulation (DBS), which are archetypical examples of “non-invasive” and “invasive” brain stimulation devices, respectively. TMS and DBS are widely used – either as an approved or as an investigational device - to achieve symptomatic treatment in a number of neurological (e.g. stroke, Alzheimer’s disease, Parkinson’s disease, epilepsy, migraine, dystonia, tremor)^4,5^, or psychiatric disorders (major depressive disorder, anxiety, obsessive-compulsive disorder)^6–9^. Although both TMS and DBS are important contemporary options in a medical team’s arsenal, they both have a number of clinical limitations. The “non-invasive” nature of TMS means that it can be applied widely in day-care clinics, and the method permits the shifting of the target within and/ or across the sessions. On the other hand, the patient is submitted to multiple repetitive sessions across days and weeks, and typically, the therapeutic benefit is transitory and requires periodic “maintenance” for it to remain or return^10^. Furthermore, TMS has low tissue penetration qualities and deep brain structures and fiber bundles cannot be targeted. The “invasive” nature of DBS permits the chronic and continuous stimulation of a selected brain structure which provides longitudinal symptom relief. On the other hand, DBS requires the precise placement of an electrode at a pre-defined and fixed target via an extensive neurosurgical procedure carried out by a specialized team. The area of the tissue activated by the implanted electrodes can be changed to some degree, but changing the target completely would entail major additional neurosurgery. In order to obtain long-term stable symptom relief and retain the option of being able to modulate activity at several targets and networks, the optimal brain stimulation technique would need to merge the targeting flexibility of TMS, with the capacity of DBS to deliver chronic stimulation precisely to any cortical or subcortical brain structure.

Progress in neuromodulation – defined as delivering better therapies for the patient - is equally dependent on improved understanding of brain networks in health and disease and on technological innovations that help to achieve previously unmet needs^11^. Ultrasound-based neuromodulation is an established stimulation modality with a unique combination of physical and biophysical characteristics: i.) it consists of a pressure wave that propagates through soft tissues with very low attenuation, requiring several centimeters of propagation to lose only half of its amplitude; ii.) it can be focused several centimeters away from the ultrasound device with moderate spatial resolution; iii.) the stimulation location can be reconfigured by means of beamforming techniques; iv.) it can safely and reversibly activate and inhibit neuronal activity without implanting the device in nervous tissue; and v.) it has built decades of public perception of safety from its use in diagnostic medical imaging^12–14^.

The current paper presents an implantable, epidural focus ultrasound device (eFUS), capable of multi-site steerable stimulation. The miniaturized focused ultrasound stimulation device consists of a two-dimensional piezoelectric transducer array directly integrated onto a custom Application-Specific Integrated Circuit (ASIC), specifically engineered for proof-of-principle neuromodulation studies in the rat brain. The system generates electronically steerable focused ultrasound with software-defined focal coordinates, achieving peak focal pressures up to 0.574 MPa (assuming an acoustic attenuation in the brain of 0.8 dB/cm/MHz), and depths of 8mm, while maintaining a sub-mm³ focal volume, sufficient to stimulate neuronal activity^15^. *In vitro* acoustic characterization confirmed accurate beam steering and focusing, while *in vivo* validation demonstrated reliable stimulation of a deep subcortical target, ventral tegmental area (VTA), a clinically relevant neuromodulation target for depression and other psychiatric disorders^9,16^. Neural activation was independently confirmed through fiber photometry recordings of dopamine release in the nucleus accumbens. Importantly, the device operated with only approximately 12 mW /MPa/DC (duty cycle) of average power consumption, generated less than 1°C of temperature increase at 10% duty cycle, remained fully functional following chronic implantation for more than 30 days, and histological analysis revealed no structural tissue damage at the deep stimulation target, collectively demonstrating the efficacy, energy efficiency, stability and safety of the technology.

## Methods and Materials

### eFUS device design, fabrication and acoustic characterization

The eFUS device comprises a 5 × 5 mm² CMOS ASIC fabricated in a 180-nm BCD process and a directly integrated two-dimensional PZT transducer array. The ASIC contains 66 × 66 transmit channels over a 4.2 × 4.2 mm² active aperture, providing 4-bit phase control for electronic beamforming and high-voltage excitation up to 20 V. An integrated temperature sensor enables monitoring of the CMOS substrate temperature during operation. For operation at approximately 4 MHz, groups of 2 × 2 ASIC channels were programmed with the same phase and connected to individual PZT elements, resulting in an approximately 33 × 33 physical transducer array. The transducer array was fabricated from a 540-µm-thick PZT5A sheet with Ni electrodes. Individual PZT pillars were defined using a DISCO DAD3240 dicing saw with a 20-µm-wide Z09-series blade. The diced array was aligned and bonded directly to the CMOS die using anisotropic conductive film (TFA22023, Telephus) and a flip-chip bonder (T-3000 Pro, Tresky). The CMOS die was subsequently wire bonded to a custom PCB, and the wire bonds were encapsulated using low-viscosity UV-curable epoxy (OG116-31, EPO-TEK). An 8-µm-thick aluminum foil, attached to the PZT surface using silver conductive paint (42469, Alfa Aesar), provided the common top ground connection. The complete assembly was coated with a 5- µm-thick parylene-C layer for electrical insulation and protection. A detailed description of the ASIC architecture, transducer integration and device characterization will be reported separately in the Results section.

### Acoustic characterization

Each eFUS device was acoustically characterized in a water tank following PZT integration and prior to implantation. Acoustic measurements were performed using a calibrated fiber-optic hydrophone (FOHSv2, Precision Acoustics; 1–20 MHz calibration range) mounted on a motorized scanning system (VK-62000, GAMPT). The hydrophone output was recorded using an oscilloscope (DSO-X 3032A, Keysight), with device operation, hydrophone positioning and data acquisition controlled through a custom graphical user interface. The operating frequency was determined individually from the acoustic response of each integrated device. Acoustic pressure was characterized at programmed focal depths of 5 and 8 mm, with the latter selected to approximate the depth of the intended in vivo targets. Two-dimensional hydrophone scans were performed to characterize the spatial profile of the focused acoustic field. Where pressure in brain tissue was estimated, the pressure measured in water was corrected assuming an acoustic attenuation coefficient of 0.8 dB/cm/MHz. A detailed description of the acoustic characterization will be reported separately in the Results section.

### Animal husbandry

Four male Sprague Dawley rats (Janvier Labs, France), aged 12 weeks were used in the study. All animals were group housed with 2-3 animals per cage until eFUS chip implantation, then single housed until perfusion (12:12 light-dark cycle; 22 ± 2 °C; 50–60% relative humidity). Animals had ad libitum access to food and water and were weighed regularly. All procedures were performed according to the regional ethics committee Regierungspräsidium Freiburg (TierSchG, G-23/080), following the EU-directive 2010/63/EU.

### Design and timeline

The experimental timeline and design are summarized in Figure 1. All animals underwent a one-week acclimatization period upon arrival at the facility and were handled prior to experiments. Surgeries were performed in two sessions to ensure both safety and optimal expression of the fiber photometry sensors. In the first stage, the viral vector encoding GRAB_DA2m_ was unilaterally injected into the left Nucleus accumbens shell. After a minimum recovery period of 3 weeks to allow for stable sensor expression, animals were implanted with an optic fiber, as well as the epidural FUS chip. The middle of the epidural focused ultrasound (eFUS) chip was positioned perpendicular to the left VTA. Following this surgery, rats underwent a minimum of 10 days of recovery and were habituated to the head-mounted implants to minimize stress and movement artifacts during the following recordings. To test the in vivo stimulation effects of the eFUS the VTA was unilaterally stimulated with different FUS parameter settings over multiple stimulation sessions over three weeks or until the hardware failed. The stimulation evoked physiological response was assessed through monitoring the dopamine release in the ipsilateral NAc using fiber photometry. Animals were sham stimulated for 30 min and then stimulated according to multiple pre-defined parameters. The rats were perfused multiple days after the last successful stimulation session and the immune response (ED1, GFAP) to the FUS stimulation and the long-term implantation of the chip was assessed histologically. Due to a limited lifetime of the chips, not all animals were tested with all parameter sets. Additionally, the alignment of the optic fiber and viral expression was checked with a GFP staining, one animal was excluded due misalignment.

**Figure 1.**
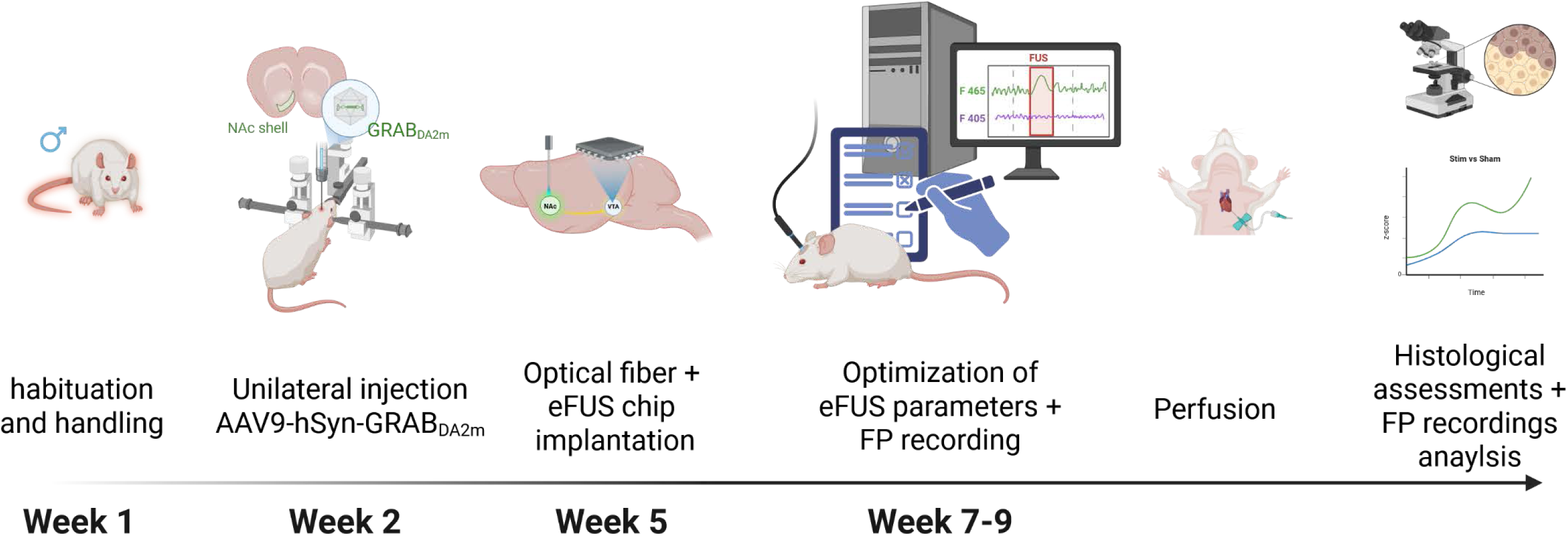
In vivo study design. Animals (6 male SD rats) underwent a one-week handling/ acclimatization period, followed by stereotactic injection of the genetically encoded dopamine sensor GRABDA2m into the shell of the nucleus accumbens (NAC). A period of at least 3 weeks was given for the expression of the sensor before the animals underwent optical fiber implantation into the NAC (for the fiber photometry measurements) and the craniotomy for the placement of the eFUS chip on the dura. A recovery period was followed by repeated stimulation and recording sessions lasting intermittently over a couple of weeks, during which different parameters were tested, and the stimulation evoked changes in dopamine release were recorded. The final session was followed by perfusion, histological analyses, and data analysis. Images created with BioRender.

### Surgery

All animals underwent two surgical procedures separated by at least 3 weeks. First, animals received Carprofen (4.0 mg/kg body weight, s.c.) for pre-emptive analgesia. Anesthesia was induced with 4% isoflurane (2l/min O2) in an induction box and maintained between 2.0-2.5 % in a stereotactic frame (Stoelting, USA). The incision site was injected subcutaneously with 5mg/kg Bupivacain for local analgesia before the initial incision. During a first surgery session animals were injected with 2x 300nl of AAV9-hSyn-GRAB_DA2m_ Dopamine sensor (Vector core, Zürich) in the left NAc shell (AP +1.2/ +1.6 mm, ML +1.1 mm, and DV −6.8 mm) at 100nl/min rate. Following surgery, animals received postoperative analgesia with buprenorphine (0.05 mg/kg, s.c.) and granisetron (0.1 mg/kg i.p.) to prevent post-surgical pica behaviour due to nausea. The days after surgery, pain was managed using Caprofen and the oral application of Metamizol (100 mg /kg; 3x d1 after surgery, then 1x per day until no more pain). In case the animal showed signs of pain additional buprenorphine doses were administered.

After at least 3 weeks of viral expression, the animals underwent a second surgical during which the optic fiber - used for the fiber photometry - and the eFUS chip were implanted (Figure 2A). An optical fiber (borosilicate, receptacle: metal ferrule MF2.5, numerical aperture: 0.66, 400 μm ⌀, Doric Lenses®, Quebec City, QC, Canada) was implanted in the left NAc shell (Figure 2B-C; AP +1.4 mm; ML +1.1 mm; DV −6.6 mm). The eFUS implantation required a craniotomy. The skull bone was carefully thinned along the edge of the planned 7.2mm x 7.2mm craniotomy and on top of the midline. To remove the skull, first the anterior edge, the midline, the posterior edge of the craniotomy was drilled without damaging the dura and the lateral edges last. Especially above the midline any small bone fragments were immediately removed to prevent bleeding from the sinus. Lastly saline was applied to the craniotomy and the edges of the two bone pieces medial and lateral from the midline were slowly detached from the dura. The dura was cleaned from any bone debris and covered with saline.

**Figure 2A-I.**
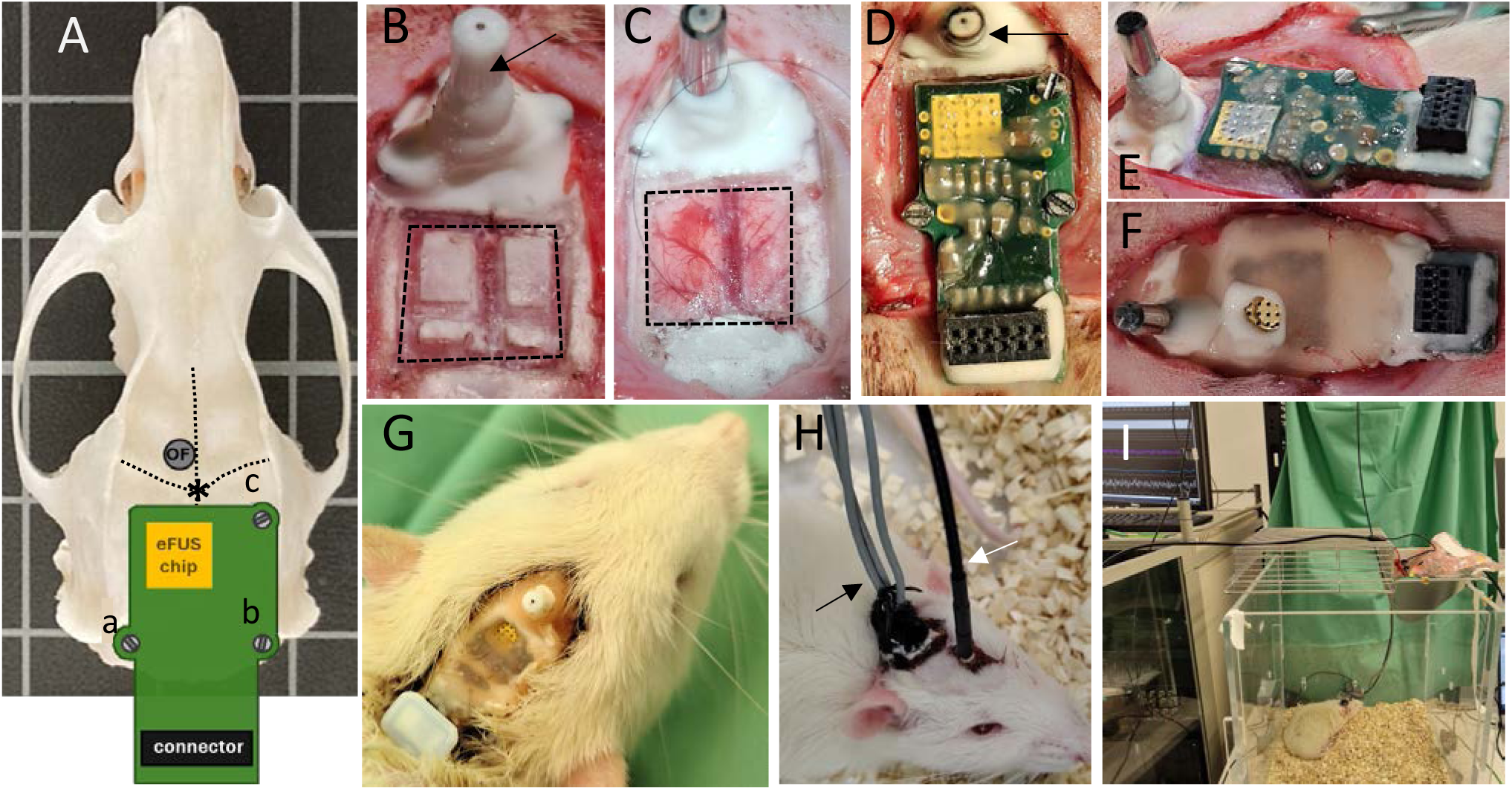
Implantation of the eFUS chip. A.) A hybrid representation of the placement of the eFUS device on the skull, with the three fixation points (a-c) of the PCB containing the eFUS chip. Further anterior from the bregma (*) is the planned implantation site of the optic fibres (OF) required for the fiber photometry monitoring of the stimulation evoked physiological changes. B-G.) The subsequent images show the craniotomy (dashed lines) and the exposed brain surface where the eFUS chip was placed. The PCB board was fixed to the skull with screws, further posterior to the optic fibres (black arrows), and the edges sealed with bone wax, skin glue and light-activated dental composite. The PCB board/ eFUS chip was also secured with dental cement to the skull. H-I.) Prior to testing, the animals were habituated to being connected up to the mother board of the stimulation (black arrow) and to fiber photometry set-ups (white arrow). Testing took place in awake and freely moving animals in a test cage.

The craniotomy placement was planned so that FUS chip could be implanted on the dura, and would have its center aligned above the target in the VTA (Figure 2B-C; AP −5.3 mm; ML +0.7, DV −8.0 mm). Based on the configuration of the eFUS chip on the PCB, the square shaped craniotomy was placed appropriately (top left corner at AP +3.2 mm ML +3.3 mm from the stimulation target), with bore holes pre-drilled at three places around the craniotomy for screws that would hold down the PCB. The eFUS chip was carefully placed on top and screwed into position. The edges of the craniotomy were sealed with bone wax, skin glue and light-activated dental composite (Figure 2D-G). The chip was additionally secured with dental cement to the bone.

### In vivo functional testing of the eFUS chips with fiber photometry recordings

After a minimum of 10 days post-surgical recovery, animals were habituated to the cables and FP patch cord. Fiber photometry recording took place in the animals’ home cage (Figure 2H-I). The in-vivo setup and the workflow are shown in Figure 3A-F. The photometric signals were recorded using FP hardware (Doric Lenses Inc., Quebec City, QC, Canada), Synapse Suite software (Version 94), and an RZ5 BioAmp processor (Tucker-Davis Technologies, Alachua, FL, USA). GFP-sensitive and isosbestic excitation signals, were generated at 465 and 405 nm, respectively. Signals were digitized at a sampling rate of 6 kHz. Transistor–transistor logic (TTL) pulses corresponding to FUS stimulation, generated by the custom-built FUS system, were transmitted to the processor and co-registered with the fluorescence signals.

**Figure 3A-F.**
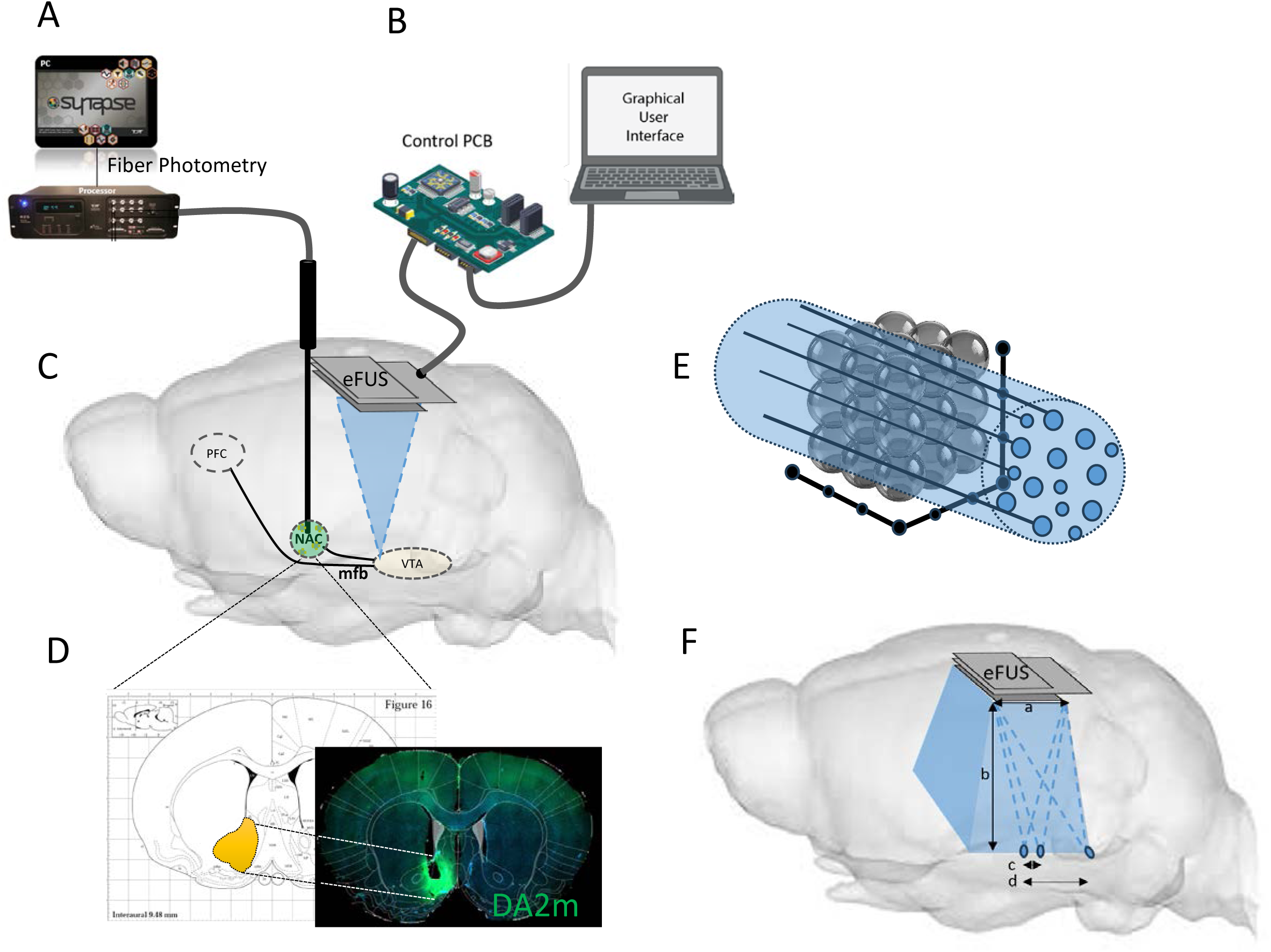
In-vivo stimulation and dopamine release monitoring. A-B.) The in vivo testing set-up integrated fibre photometry, to monitor and record changes in dopamine release, and eFUS, including the control PCB/ motherboard and a graphical user interface for the spatial and temporal control of the stimulation. C.) The optic fibre was implanted into the nucleus accumbens (NAC) to monitor stimulation evoked changes in dopamine release as a consequence of focus ultrasound stimulation of the anterior part of the ventral tegmental area (VTA), including the midbrain dopaminergic projections coursing through the medial forebrain bundle (mfb) to the NAC and the prefrontal cortex (PFC). D.) Post-mortem analysis confirmed the correct placement of the biosensor sensitive to changes in dopamine (GRAB_DA2m_) levels in the NAC. E.) Schematic diagram demonstrating the steerability capacity of the eFUS to project the focal point in 3D. During the testing targeting the anterior part of the VTA/ mfb, the focal point could be displaced by 0.5mm in the z-axis, and 0.5 mm in the xy-axis. This was done in order to identify the optimal “spot” with the strongest physiological response. F.) In theory, the eFUS device has a range that depends on chip’s technical parameters such as its aperture (a), maximal focal depth (b), and the minimal (c) and maximal steering (d) capacity.

Animals were stimulated with twelve different sets of FUS parameters over multiple recording sessions (Table 1) to test the influence of duty cycle (DC) and sonic duration (SD) with similar total stimulation times, as well as pulse repetition frequencies (PRF). Fiber photometry recordings measuring changes in dopamine release was continuous, but the baseline, the stimulation, and the post-stimulation data used for the analysis is summarized in Figure 4. Each stimulation parameter condition was repeated 3 times. Since the focal beam could be steered up to a depth of 10 mm, the stimulation with each set of parameters was also repeated at different depths (AP −7 to −9mm) to ensure best coverage of the target area. During stimulation, the real-time DA release in the NAc was measured using fiber photometry. For each animal only the data from the most promising stimulation location was selected for further analysis.

**Figure 4.**
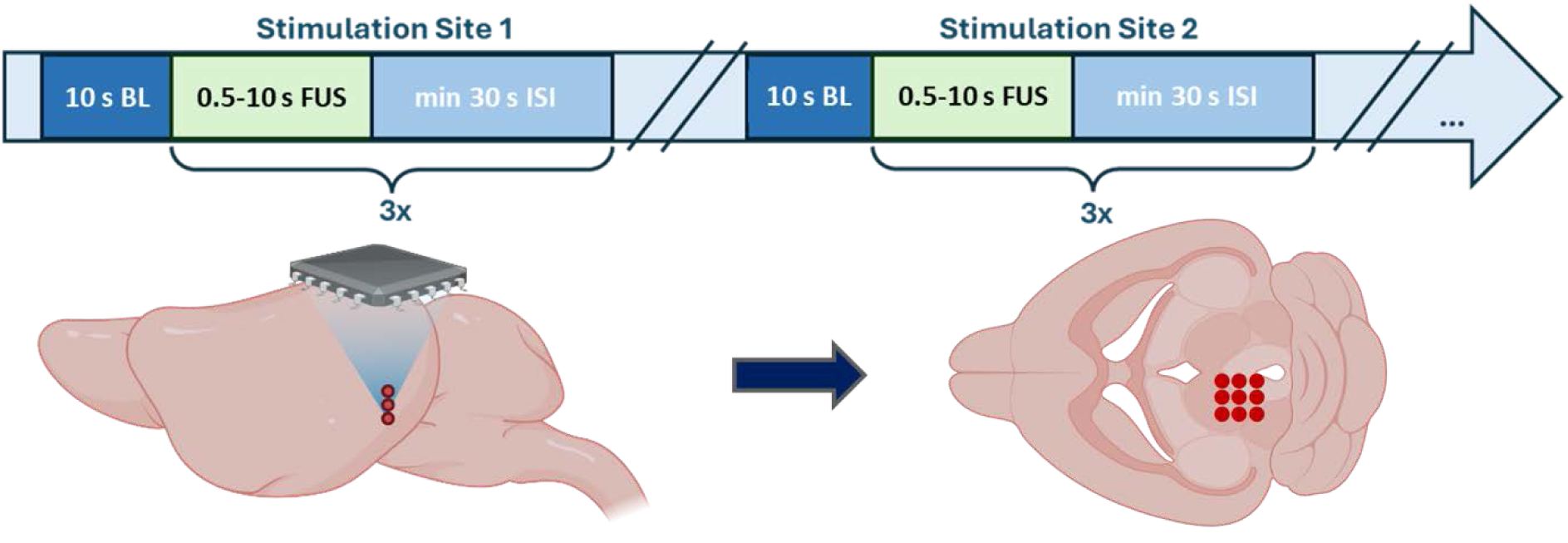
Protocol design of the recorded and analysed data. Fiber photometer recording was continuous. Multiple stimulation parameters were tested and each parameter was repeated 3 times. The analysed sessions were composed of baseline recordings (BL, 10s), a stimulation period (FUS, lasting between 0.5-10s depending on the parameter), and a post-stimulation or inter-stimulation interval (ISI, min 30s). Including the change of parameters and testing the parameters at different sites, the testing session lasted approximately 30 mins per animal.

**Table 1.** Summary of the twelve FUS parameter combinations (p1-p12) evaluated for the optimization of dopamine release. Parameters p1-p5 (highlighted) were the combination sets that were analyzed in more detail in the Results section. In all cases, the stimulation voltage was maintained at 20 V, the baseline duration was 30s, and the post-stimulation recovery period was 45s. Sonication (stimulation) duration ranged from 0.05s to 10s. Peak pressure depended on the individual chip and ranged between 0.34 and 0.57 MPa across all sets. *p6 was rarely tested set due to very short stimulation time (SD).

| Parameter combination | Pulse Width (PW; μs) | Pulse Repetition Frequency (PRF; kHz) | Duty Cycle (DC; %) | Sonication Duration (SD; s) | Inter-Stimulus Interval (ISI; s) |
| --- | --- | --- | --- | --- | --- |
| p1 | 25 | 1 | 2.5 | 10 | 32.1 |
| p2 | 50 | 1 | 5 | 5 | 31.1 |
| p3 | 100 | 1 | 10 | 2.5 | 33.6 |
| p4 | 200 | 1 | 20 | 1 | 35.1 |
| p5 | 500 | 1 | 50 | 0.5 | 35.6 |
| p6* | 300 | 1.5 | 45 | 0.05 | 36.05 |
| p7 | 40 | 1.5 | 6 | 0.2 | 35.9 |
| p8 | 200 | 0.2 | 4 | 1 | 35.1 |
| p9 | 3846 | 0.13 | 50 | 0.5 | 35.6 |
| p10 | 231 | 0.13 | 3 | 0.5 | 35.6 |
| p11 | $10 \times 10^3$ | $2.5 \times 10^{-3}$ | 2.5 | 5 | 31.1 |
| p12 | $80 \times 10^3$ | $2.5 \times 10^{-3}$ | 20 | 1 | 35.1 |

The eFUS chips utilized in this study were designed for a fundamental frequency of 4 MHz, but individual characterization showed peak pressures varying between 3.7 and 4.0 MHz. Similarly, the peak pressure measured in vitro with a hydrophone at 8 mm distance varied between devices.

For comparison, all animals also underwent a 30 min sham stimulation session which corresponded to the minimum length of normal stimulation session. During these sessions the animals were connected up but no stimulation was applied. There was no correction for a possible auditory confounder although it was assumed the epidural nature of the stimulation would minimize bone conductance and therefore auditory side effects that have been associated with clinical transcranial FUS^17,18^.

### Histology

Animals were transcardially perfused multiple days after the last stimulation session with PBS and 4 % PFA. Brains were postfixed overnight and transferred to 30% sucrose solution at 4°C until further processing. Brains were cut with a slide microtome into 40µm coronal sections. For the free-floating DAB staining sections were blocked for 2h in 5% BSA solution and stained with α-CD68 (ED1) (mouse, 1:300, Millipore) and α-GFAP (rabbit, 1:600, Dako) overnight. The biotinylated secondary antibodies (anti-mouse IgG-biotinylated; goat; 1:200; Vector Laboratories and anti-rabbit IgG-biotinylated; goat; 1:200; Dako) were applied for 1h at RT and the reaction was visualized with a DAB peroxidase substrate Kit (Vector Labs). To check for the accuracy of the viral injection and optic fiber placement, brains were additionally fluorescent stained with anti-GFP antibody (mouse, 1:500; Invitrogen) over night and the anti-mouse-A488 (goat; 1:500; Life Technologies) secondary antibody, as well as DAPI (1:200; Sigma Aldrich).

### Fiber photometry analysis

The fiber photometry data from each stimulation site was extracted using pMAT v1.2 (The Barker Lab, Philadelphia, PA, USA) and a custom-made MATLAB R2025b (The MathWorks Inc., Natick, MA, USA) script. The raw GFP signal (465 nm, F_465_) from the GRAB_DA2m_ dopamine sensor activity was fitted to the isosbestic signal (405 nm, F_405_) in order to obtain the change in corresponding dopamine fluorescence (*ΔF/F* = *F*_465_−*F*_405_/*F*_405_) using pMAT. The ΔF/F signal was then further processed using custom scripts. The 10s pre stimulation as well as 30 s post stimulation start were extracted for each stimulation repetition and saved in separate files according to stimulation location.

In a second step, only the sham data and the optimal stimulation location for each animal were further processed. The data was normalized to the event related baseline (10 s pre-stimulation). This results in the robust event related z-score:

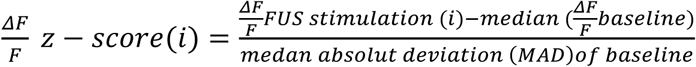

The three repetitions for the stimulation with each parameter set were averaged within each animal. In case a certain parameter set was repeatedly used the stimulations were averaged.

The area under the curve (AUC) during stimulation and up to 10 s post stimulation was calculated as well as the peak amplitude post stimulation onset. The AUC represents the total cumulative change in fluorescence signal intensity over a specific time window, and it is a reflection of net magnitude of neurotransmitter release. All calculated values and z-scores were then averaged across animals. The z-score was taken to represent the normalized fluctuation in fluorescence intensity over time, calculated by subtracting the baseline mean and dividing by the standard deviation.

### Assessing targeting using microbubbles

Targeting accuracy was initially assessed prior to the recordings in two separate animals by opening the blood–brain barrier (BBB) using microbubbles. The eFUS chip was implanted under anaesthesia, and 5 × 10^8^ SonoVue © microbubbles, as well as 4–5% Evans Blue dye, was injected into the tail vein of the animal. Several targets at various depth were stimulated using the same set of parameters for up to 2 minutes. Through BBB opening at the focal spot, the dye was able to extravasate into the surrounding tissue. The animals were sacrificed 2 hours later and prepared for fluorescence microscopy to detect even small amounts of dye.

### Statistics

The area under the curve (AUC) during the stimulation (0.5-10 s) and 5 s post-stimulation, as well as the peak z-score during and 5 s post stimulation were extracted for each animal’s sham and stim conditions by the custom Matlab script. The statistical analysis as well as the graphs were generated using GraphPad Prism 9.5.1. Normal distribution of data was assessed through a QQ-plot. A paired t-test was performed to compare the Stim vs Sham data for each set of parameters.

## Results

### Technical characterization of the eFUS device

Electrical characterization confirmed the operation and programmability of the integrated transmit electronics (Figure 5A). The 4-bit phase control provided 16 programmable phase states across the transmit channels, enabling the relative timing of the excitation signals to be configured for electronic focusing and steering. The high-voltage output stages generated the programmed excitation waveforms up to a 20 V supply, while the output voltage and duty cycle could be independently configured according to the stimulation protocol. Following electrical characterization, the PZT array was directly integrated onto the CMOS ASIC and the assembled device was wirebonded and packaged on a custom PCB (Figure 6B). Acoustic characterization in water confirmed the formation and electronic steering of a spatially confined focus (Figure 5C–D). At 4 MHz, a 5 V drive and a programmed focal depth of 5 mm, the measured acoustic field exhibited a lateral full-width at half-maximum (FWHM) of approximately 300 µm and an axial depth of field of approximately 2.4 mm. Electronic steering of the focus was demonstrated by applying different phase configurations across the array, producing the expected lateral displacement of the acoustic focus. Focusing was also demonstrated at larger programmed depths (8 and 10 mm), covering the range required for the subsequent in vivo experiments. Acoustic output was characterized individually for six integrated devices to quantify device-to-device variation following fabrication (Figure 5E). After accounting for an acoustic attenuation of 0.8 dB/cm/MHz in brain tissue, the estimated peak pressure was 0.56 ± 0.16 MPa at 5 mm and 0.41 ± 0.12 MPa at 8 mm (mean ± SD, n = 6), with individual 8-mm values ranging from 0.26 to 0.57 MPa. Thermal characterization was performed under pulsed operation at 4 MHz and 20 V to assess heating of the device during stimulation (Figure 5F). The temperature increase remained below 1 °C at 1% duty cycle and reached approximately 1 °C at 10% duty cycle, whereas operation at 50% duty cycle resulted in substantially greater heating, with a temperature increase of approximately 6 °C by the end of the stimulation period.

**Figure 5A-F.**
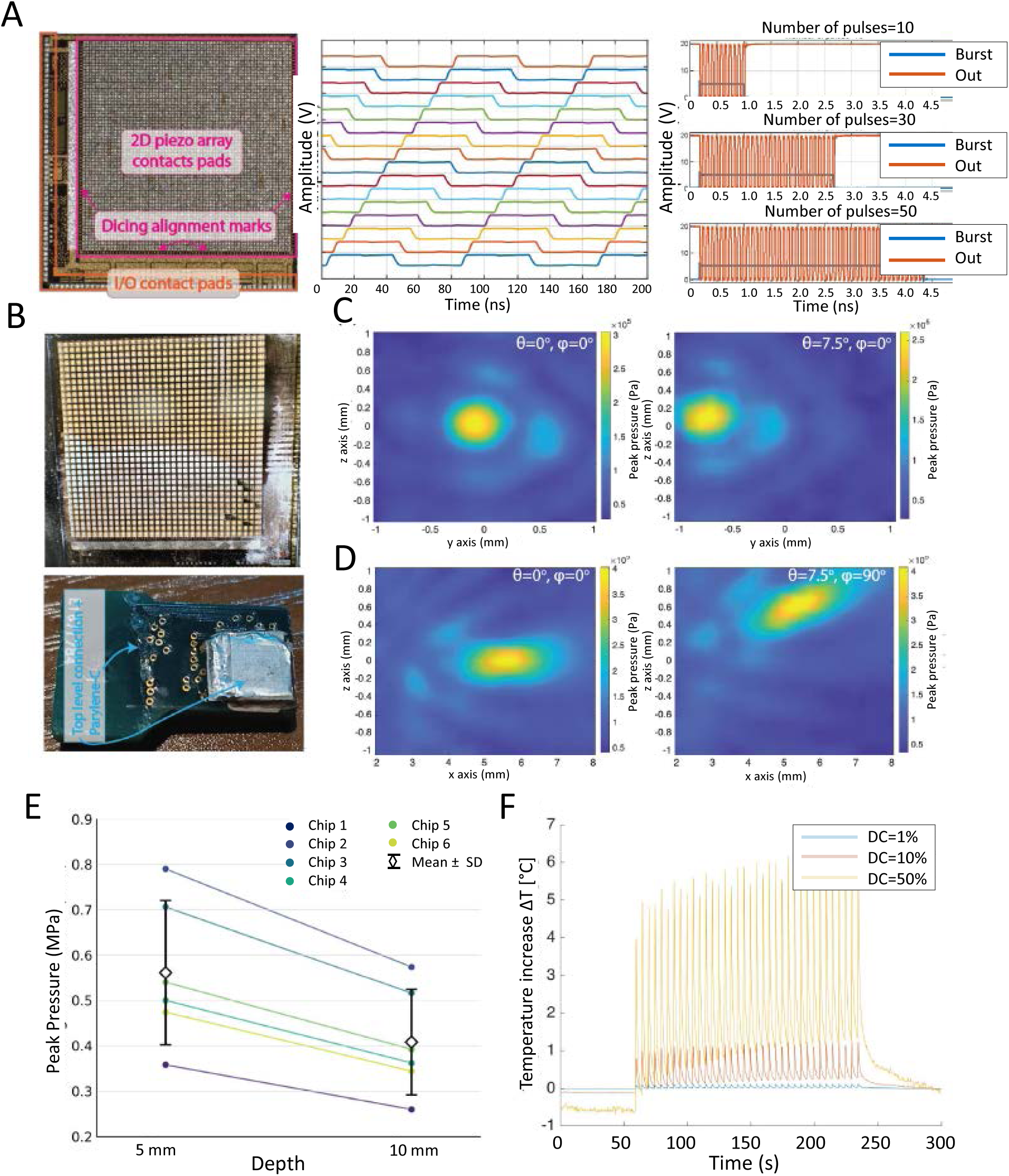
Technical characterization of the eFUS device. A.) CMOS ASIC and electrical characterization. 5 mm x 5 mm die micrograph showing the 4.2 × 4.2 mm² two-dimensional transducer interface, together with measured transmit waveforms demonstrating programmable phase control with eight different phases, and burst generation with different burst durations. B.) Integration of the diced PZT array onto the CMOS ASIC and photograph of the assembled eFUS device following packaging, top-electrode connection and final encapsulation with a layer of Parylene-C. C.) Measured transverse acoustic pressure distributions in water for a programmed focal depth of 5 mm, showing the non-steered focus (θ = 0°, φ = 0°) and electronic steering by θ = 7.5° along φ = 0°. D.) Corresponding axial acoustic pressure distributions showing the non-steered focus and electronic steering by θ = 7.5° along φ = 90°. Acoustic field measurements in (C,D) were performed at 4 MHz with a 5 V drive voltage. E.) Device-to-device acoustic output for the six eFUS devices characterized at focal depths of 5 and 8 mm, all driven at 20 V. The optimal frequency was chosen beforehand with a frequency sweep, with devices 1 and 2 working at 3.69 MHz, and devices 3-6 at 3.76 MHz. Connected points represent measurements from the same device and black markers and error bars indicate mean ± SD (n = 6). Pressures correspond to estimates in brain tissue derived from water-tank measurements using an acoustic attenuation coefficient of 0.8 dB/cm/MHz. F.) Temperature increase measured at the device during pulsed operation at 4 MHz and 20 V for duty cycles of 1%, 10% and 50%. Measurements were performed with a pulse repetition frequency of 1 kHz, 0.5-s sonication duration and 4.5-s inter-stimulation interval, comprising 1 min baseline, 3 min active operation and 1 min post-stimulation recording.

**Figure 6A-C.**
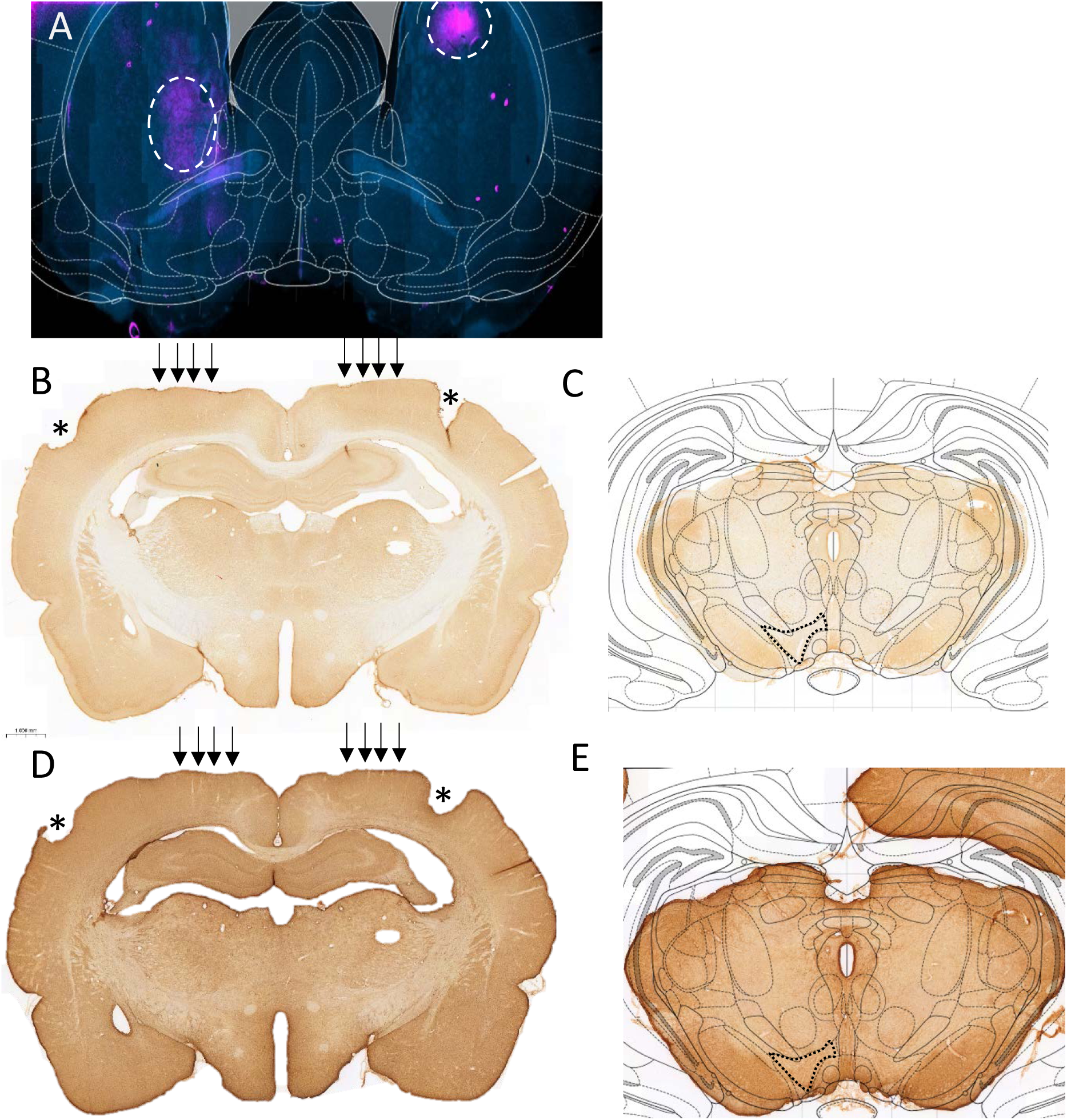
Post-mortem assessment of targeting and safety issues. A.) Animals received tail vein injection of microbubbles diluted in 4–5% Evans Blue dye and various predetermined targets were stimulated with the implanted eFUS. Through blood-brain-barrier opening at the focal spot, the dye was able to extravasate into the surrounding tissue (white dashed line). B and D.) Histological assessment identified minor tissue impact associated with the implantation of the eFUS device, manifested as cortical damage induced by the fixation screws (*) and the flattening of the cortical surface in contact with the piezoelectric transducer array (black arrows). Inflammatory markers ED1 (B) and GFAP (D) showed little or no activity. C and E.) The regions in the proximity of the target area (black dotted lines), at the junction of the anterior part of the ventral tegmental area and the medial forebrain bundle, were absent of ED1 (C) and GFAP (E) expressing cells, indicating the absence of tissue damage, and safety of the stimulation.

### Histology

Acoustic cavitation, produced by microbubbles interacting with focused ultrasound, can open the BBB and permit a dye to extravasate into the tissue. This method was used to confirm that the eFUS chip emitted sufficient focal pressure at a targeted area. Likely due to the limited half-life of microbubbles and the decrease in focal pressure when the ultrasound beam is steered, the strongest signal was consistently observed at the stimulation location perpendicular to the center of the chip. Fluorescence microscopy confirmed Evans Blue extravasation, suggesting that perpendicular stimulation is mostly spatially accurate in all axes (Figure 6A). This also indicates that the stereotactic implantation was precise and the chip was not tilted. On the contralateral hemisphere, the focal spot appeared either rather broad or concentrated around a blood vessel. In both cases, the BBB opening appeared to be slightly more lateral or ventral than intended. These results suggest that perpendicular stimulation is accurate, and that the chip’s steering capability remains useful for identifying the stimulation location producing the strongest effect within the target region.

To assess possible implantation and stimulation-associated inflammatory responses, indicated by the presence of macrophages/ activated microglia or reactive astrocytes, ED1 and GFAP, respectively, immunohistochemistry was performed on coronal brain sections obtained after completion of the experimental procedures. Overall, both ED1 and GFAP immunoreactivity were low to moderate across the experimental cohort and was predominantly localized to regions associated with implant trajectories of the optic fiber used for fiber photometry. Some consequences related to the surgical procedures required to implant the eFUS were observed as superficial cortical damage caused by the screws fixing the eFUS chip to the skull (Figure 6B and D; ED1 and GFAP, respectively), but no sign of tissue damage/ inflammation was observed at the deep brain structure, the ventral tegmental area, that was targeted by the focused ultrasound stimulation (Figure 6C and E; ED1 and GFAP, respectively).

### In vivo functional testing

A total of twelve different sets of stimulation parameters were tested to identify the combination most effective for inducing dopamine release. The parameters varied in pulse width (PW; 25 µs to 80.000 µs), pulse repetition frequency (PRF; 0.0025 kHz to 1.5 kHz), and duty cycle (DC; 2.5% to 50%), while the voltage was kept constant at 20 V and the peak pressure ranged between 0.26 and 0.574 MPa. Five parameter combinations (p1, PRF 1kHz, DC 2.5%, SD 10s; p2, PRF 1kHz, DC 5%, SD 5.0s; p3, PRF 1kHz, DC 10%, SD 2.5s; p4, PRF 1kHz, DC 20%, SD 1.0s; p5, PRF 1kHz, DC 50%, SD 0.5s) have been selected for detailed analysis. The baseline duration was set to 30s min for all trials, and post-stimulation recovery period was 40s. The stimulation paradigm consisted of a sequenced application of ultrasound bursts. Each trial began with a 30-second baseline (BL) recording (only last 10 s used for analysis), followed by a sonication duration (SD) of 0.5 to 10s and concluded with an inter-stimulus interval (ISI) of at least 45 s. The sequence of SD and ISI after each baseline was repeated 3 times per stimulation site to ensure the stability and reproducibility, but also look for a trend of the dopamine response. For each individual parameter set, sonication was systematically delivered at varying tissue depths. The most common locations evaluated along the dorsoventral axis were at depths of 7.0, 7.5, 8.0, 8.5, and 9.0 mm. Depending on where the most robust functional outcomes were achieved along this axis, multiple anteroposterior (AP) and mediolateral (ML) coordinates were subsequently evaluated to precisely identify the optimal stimulation target for modulating the dopaminergic signaling.

Consistent with the targeting accuracy confirmed through BBB opening in separate animals, the most robust results were achieved at 8 mm depth with no medio-lateral adjustment. Only one animal showed more consistent results at 8.5 mm stimulation depth.

Figure 7A-E summarizes the results from the five different stimulation parameter combinations. The average z-scores for the parameter settings p1-5 over the baseline period (−10 s before stim onset) up to 30 s post-stimulation onset are shown in Figure 7A. The z-score represents the normalized fluctuation in fluorescence intensity over time, calculated by subtracting the baseline mean and dividing by the standard deviation. None of the tested parameters evoked a sharp onset of dopamine release during the stimulation period (marked in red in Figure 7A, or summarized in Figures 7B and 7C). Dopamine activity did not increase significantly over baseline levels with parameters p1 and p2, which included lower duty cycles (2.5 and 5 %, respectively). However, for p3-5, an increase in dopamine can be seen post-stimulation, with a peak latency of ca. 6.5s (p3, p4) to 12s (p5) after stimulation offset. p3 with 10 % DC and 2.5 s of sonication duration, showed a significantly higher AUC (Figure 7D; paired t-test: t(2)=6.32, p=0.024) and peak z-score (t(2)=8.16, p=0.015) in the 5s post stimulation compared to sham conditions. Similarly, p5 likewise exhibited a significant increase in dopamine levels; however, this finding should be interpreted with caution given the limited sample size (Figure 7D; n = 2; paired t test AUC 5 s post stim t(1)=50.71, p=0.013; Figure 7E; peak z-score t(1)=15.57, p=0.041). p4 also demonstrated a small increase in dopamine levels around 6.5s post stimulation onset, although the effect did not reach significance. Notably, both p4 and p5 exhibit dopamine responses characterized by shorter duration and reduced peak amplitude compared with p3, which remained above sham levels ca. 20 s post stimulation (Figure 7E). No significant correlation was observed between peak pressure and effect size for p4, although a negative linear trend was apparent, with lower responses observed at increasing pressures. However, this relationship may have been influenced by differences in target engagement, particularly the positioning of the focal spot. Given the limited sample size, further experiments with a larger number of animals will be required to determine whether this trend represents a consistent relationship between stimulation pressure and response magnitude.

**Figure 7A-E.**
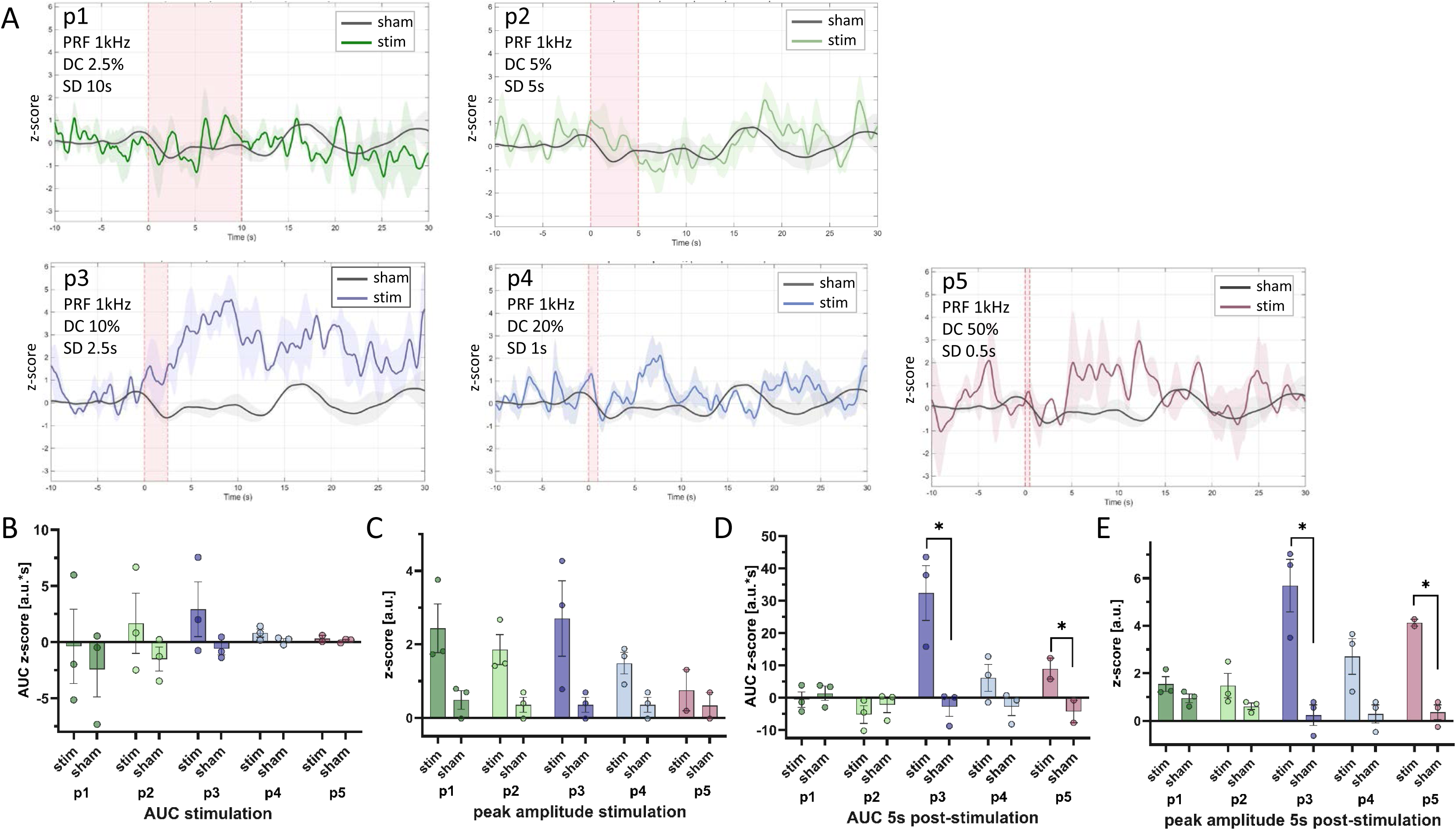
Dopamine release in the NAc in response to unilateral eFUS stimulation of the VTA using five different stimulation parameter sets (p1–5). A.) Mean z-score during the baseline period (−10 s to stimulation onset) and 30 s following stimulation onset for five stimulation parameter sets (p1–5). Sonication duration is indicated in red. Responses represent the mean of three repetitions per animal (n = 3 for p1–4; n = 2 for p5). Sham, mean response during a 30-min recording without FUS or auditory masking. B.) AUC during sonication for stimulation and sham conditions. C.) Peak z-score amplitude during stimulation. D.) AUC during the 5 s following stimulation. Significant differences between stimulation and sham conditions were observed for p3 and p5 (paired t-test). E.) Peak z-score amplitude during the 5 s following stimulation; significant increases were observed for p3 and p5 (paired t-test).

## Discussion

The study provides the first *in vivo* physiological assessment of a novel, innovative and implantable epidural focus ultrasound (eFUS) device capable of steerable neurostimulation of deep, subcortical brain structures. The miniaturized eFUS device consists of a two-dimensional piezoelectric transducer array directly integrated onto a custom ASIC, specifically engineered for proof-of-principle neuromodulation studies in the rat brain. *In vitro* acoustic characterization confirmed accurate beam steering and focusing capabilities, while the current *in vivo* application tested its ability to stimulate a deep subcortical target, the ventral tegmental area (VTA), a clinically relevant neuromodulation target for depression and other psychiatric disorders. Neural activation was confirmed through fibre photometry recordings of dopamine release in the nucleus accumbens.

To our knowledge, the eFUS platform represents the first fully integrated two-dimensional phased-array ultrasound system enabling three-dimensional electronic beam steering for chronic neuromodulation in freely moving animals, without significant thermal accumulation under the stimulation conditions used in vivo. Compared with the integrated phased-array system reported elsewhere^19^, the present device generates a 3.3-fold higher peak-to-peak focal pressure (3.8 versus 1.15 MPa) for the same focal depth, despite using a threefold lower maximum driving voltage (20 versus 60 V), corresponding to an approximately 9.9-fold improvement in focal pressure per volt (190 versus 19.2 kPa/V). Maximum power consumption at 100% duty cycle is reduced by approximately 12.7-fold, from 19.2 to 1.514 W. Under pulsed operation, the measured temperature increase was approximately 1 °C at 10% duty cycle, compared with 79 °C at 7.5% duty cycle reported for the previous system. Relative to our earlier PZT-on-CMOS implementation^20^, the present device increases the focal peak-to-peak pressure at 5 mm by 38-fold, with an approximately 9.5-fold higher focal pressure per volt. These advances in integration density, acoustic output and power efficiency enable electronic control of a spatially confined ultrasound focus at the depths and pressure levels required for the chronic in vivo experiments reported here.

Mechanism of DBS, and how acute and chronic DBS of midbrain structures and projections impact on dopamine and noradrenalin release in the nucleus accumbens and prefrontal cortical regions, have been thoroughly investigated in in vivo freely moving animals using fibre photometry^21–24^. The clinical relevance – particularly in the context of experimental therapies for Treatment Resistant Depression - of targeting midbrain and medial forebrain bundle structures using neurostimulation modalities have been described elsewhere ^16,25–27^. The current work was designed as an explorative, proof-of-principle study validating the surgical and in vivo protocols associated with the novel, miniaturized implantable eFUS device. It focused on testing – in awake and freely moving animals - a diverse range of stimulation parameters with the objective of identifying specific eFUS conditions that evoked specific stimulation-mediated physiological responses. Specifically, the device was used to target the mesolimbic projections originating from the ventral tegmental area and projecting to forebrain regions and integrated fiber photometry method was applied to monitor stimulation evoked dopamine release in the nucleus accumbens.

The stimulation frequency used in the current study was engineered to be 4MHz, and could not be altered. However, the parameter combinations selected and analyzed in more depth were designed to evaluate the physiological impact of an increasing duty cycle (DC; scaling from 2.5% to 20%) paired with a corresponding reduction in total sonication duration (SD; decreasing from 10 s to 1 s).

The fundamental frequency (FF) of the tested device (4 MHz) is substantially higher than that commonly used for transcranial focused ultrasound neuromodulation in clinical trials (typically ∼250–650 kHz)^28^. The higher frequency provides a more precise focal spot and can therefore be advantageous for targeting small structures. But since outside of epidural FUS the clinical relevance remains low due to the high skull attenuation the effects of fundamental frequency on the physiological stimulation response are currently not well researched.

The increased spatial precision afforded by the higher carrier frequency also places greater demands on targeting accuracy when stimulating small structures. To account for this, different target locations were tested. The high accuracy of targeting demonstrated by BBB opening provides evidence that the intended focal region can be reached reproducibly; however, the current sample size is insufficient to determine whether variations in peak pressure or focal positioning are systematically related to response magnitude.

In addition to the carrier frequency, the pulse repetition frequency (PRF) may be an important determinant of the neuromodulatory response. Increasing evidence suggests that the temporal structure of the acoustic stimulus, including PRF, can influence the magnitude and direction of the neural response. In particular, PRF may modulate the tissue displacement dynamics associated with acoustic radiation force especially for higher FFs^29^. Which is why, in the present study, the PRF was fixed at 1 kHz, thereby restricting the parameter space explored at the relatively high end of the commonly investigated PRF range. There is also some preliminary evidence that higher PRFs could more likely evoke an excitatory response^30,31^. By maintaining a constant pulse repetition frequency (PRF) of 1 kHz, this approach kept the net active ultrasound exposure time identical at 0.25 s across the conditions, thereby isolating the effects of temporal energy distribution rather than cumulative energy delivery.

As this was an exploratory study aimed primarily at establishing the capacity of the device to elicit a biological response, the maximum available acoustic pressure was used, while the five parameter combinations (p1-p5) selected and analysed in more depth were designed to evaluate the physiological impact of an increasing duty cycle (DC; scaling from 2.5% to 50%) paired with a corresponding reduction in total sonication duration (SD; decreasing from 10 s to 0.5 s). The results showed no indication of a dopamine release in response to stimulation with lower DC (2.5, 5 %), while the 10 % DC (p3) elicited the strongest increase in dopamine post stimulation, with both the duration of transmitter release and the peak amplitude being significantly higher compared to the sham stimulation condition. The absence of a measurable response at the lower duty cycles tested could be interpreted in light of one potential mechanism of action of FUS, the neuronal intramembrane cavitation excitation (NICE), which suggests T-type calcium currents contribute to charge accumulation during the OFF time and thereby favor activation of low-threshold-spiking interneurons under specific low-duty-cycle stimulation regimes. This mechanism has been proposed to account for inhibitory network effects at low duty cycles and excitatory effects at higher duty cycles^32,33^. However, although the stimulation conditions p4 and p5 with 20 % and 50 % DC, respectively, did show an increase in dopamine post stimulation which was significant in case of p5, the effect was shorter and less pronounced than under 10% DC. This could suggest that while higher duty cycles are favorable, a minimum sonication duration might be needed to see effects independent of the total ON time. Our results show that the eFUS device was capable of eliciting a significant, parameter dependent increase in dopamine when stimulating the VTA, validating the device’s capability to modulate deep targets.

The proposed mechanisms of action of focused ultrasound mediated neuromodulation, clinically referred to as low intensity (LI) FUS, are numerous and include modulation of action potentials - both excitatory and inhibitory-via electrophysiological-mechanical coupling, modulation of mechanosensitive ion channels, or cavitation and sonoporation^12,34^. Besides directly modifying neuronal and network activity, LI-FUS is also associated with neuroprotective effects, both by permitting drug delivery via opening of the BBB, or via the enhanced local release of neurotrophic factors^35^. To date, most of pre-clinical and clinical testing of LI-FUS occur using transcranial application of the transducer, and there are a large number of publications in both animals and humans confirming the neuromodulatory capability of tFUS^13^. An experimental exception to transcranial application was the study using an implantable ultrasound stimulator for deep brain activation, but this approach remains spatially restricted similar to DBS^36^.

Clinical trials assessing the therapeutic effects of tFUS are in the majority in the context of neurological patients, but the approach is also being considered with psychiatric patients ^37–40^. However, all clinical trials, whether in neurological or psychiatric disorder, are using tFUS, and at this moment none are looking at implantable devices similar in concept to the one tested in the current study. tFUS can reach diverse deep brain structures, but stimulation can only be done in specialized clinics, with typically repetitive daily stimulation lasting between minutes to a few hours maximum. Experience from DBS and TMS in Parkinson’s disease, Obsessive-Compulsive Disorder and Major Depressive Disorder suggests that long-term and stable symptom relief requires continuous and chronic stimulation^10,41,42^. Furthermore, tFUS is impacted by skull thickness and low Skull Density Ratio is associated with reduced treatment efficacy^43,44^. Skull thickness would be irrelevant for any epidurally placed LI-FUS device, which would be capable of delivering therapeutic stimulation both intermittently or chronically, according to requirements.

Multiple valid stimulation targets exist in neurological and psychiatric disorder, and the medical team need to choose one from several other potential ones. The choice of target is typically made according to the patient’s dominant symptom (for example in Parkinson’s’ disease^45^) or other factors, including the medical team’s own ideas concerning the key neural hub responsible for the disorder (as in for example is the case in MDD^46,47^). Single and consensual stimulation targets are rare. A key innovation and novelty of the proposed epidural focused ultrasound stimulation technology would be that it proposes both high spatial resolution and steerability, permitting a large degree of freedom targeting a variety of currently recognized therapeutic neuronal hubs in both in cortical and deep-seated brain structures. Stimulated targets could be re-programmed - based on integrated cortical recording capacity, or manifested symptoms - without re-operation, keeping the approach minimally invasive while adaptive even to evolving symptoms.

Several methodological considerations should be taken into account when interpreting the present findings. From a technical perspective, the acoustic field of each eFUS device was characterized in water prior to implantation, while the pressure at the targeted brain location could not be measured directly in vivo. The reported in vivo pressures therefore represent estimates based on the water-tank measurements and an assumed brain attenuation coefficient of 0.8 dB/cm/MHz. Propagation through the dura and brain tissue, as well as variations in acoustic coupling and device positioning after implantation, may alter the pressure and spatial characteristics of the focus relative to the hydrophone measurements. Although epidural placement through a craniotomy avoids transmission through the skull over the active aperture, the acoustic field was not mapped after implantation. Device-to-device variation in acoustic output was also observed following PZT integration; this was addressed by characterizing each device individually before implantation, but resulted in differences in the estimated pressure delivered across animals. Temperature was monitored at the device level during characterization, but local temperature changes at the acoustic focus were not measured in vivo. In the present device, the maximum usable duty cycle is ultimately constrained by heat dissipation at the chip–dura interface, limiting the range of acoustic stimulation parameters that can be explored. Further reductions in electrical power consumption and improved thermal management will therefore be required to provide greater flexibility in duty cycle and sonication duration. The maximum transducer driving voltage was also limited to 20 V. Higher acoustic pressures may require increased driving voltages in future implementations, although this will need to be balanced against the associated increase in electrical power dissipation and heating at the device–tissue interface. Furthermore, the sham condition did not include auditory masking or otherwise control for potential auditory confounds associated with stimulation onset and offset^48^. Although previous clinical studies suggest that the predominant effect is mediated by ultrasonic vibrations conducted through the skull to the inner ear, a contribution of auditory stimulation cannot be fully excluded^17,18^. Future experiments incorporating an appropriate auditory control condition will therefore be important to confirm the specificity of the observed response to FUS stimulation. Finally, the relatively small sample size, particularly for some parameter conditions, limits the statistical power of the study and warrants confirmation of the present findings in a larger cohort.

## Conclusion

A novel and innovative miniaturized eFUS device, consisting of a two-dimensional piezoelectric transducer array, was tested both in vitro and in vivo. Compared to the state of the art, the device combines approximately one order of magnitude higher acoustic efficiency (190 kPa/V), one order of magnitude lower average power consumption (12.25 mW/MPa/DC), and nearly two orders of magnitude lower self-heating (<1°C at 10% duty cycle). The device was capable of delivering focal pressures exceeding 1.9 MPa, which is above threshold level for neurostimulation, at depths of 5 mm while maintaining a sub-mm³ focal volume. The preliminary in vivo data described in the paper supports biological compatibility of the device, steerability of the focal point, and the possibility to perform chronic functional testing, with repetitive stimulation across diverse stimulation parameter conditions. Neuromodulation capacity targeting deep brain structures of the eFUS chip was also validated by demonstrating its capacity to modulate dopamine release in awake and freely moving rats following stimulation of the ventral tegmental area. The device proposes a novel and innovative approach to neuromodulation, with more flexibility and adaptability to select stimulation targets customized to the patients’ evolving needs, compared to current available technologies.

## Funding sources

This work was supported by the Dept. of Stereotactic and Functional Neurosurgery, Clinic, Medical Center - University of Freiburg (Germany), and by the UPSIDE project that has received funding from the European Union’s Horizon Europe EIC-PATHFINDER program under grant agreement No 101070931.

## Declaration of generative AI in scientific writing

During the preparation of this work the authors did not use any AI tools and take full responsibility for the content of the published article.

## Conflict of interests

The authors declare no financial or personal conflicts of interest.

## Notes

### Competing Interest Statement

The authors have declared no competing interest.

